# Exposure to Zika and chikungunya viruses impacts aspects of the vectorial capacity of *Aedes aegypti* and *Culex quinquefasciatus*

**DOI:** 10.1101/2023.02.13.526953

**Authors:** Mônica Crespo, Duschinka Guedes, Marcelo Paiva, Mariana Sobral, Elisama Helvecio, Rafael Alves, George Tadeu, Claudia Oliveira, Maria Alice Varjal Melo-Santos, Rosângela Barbosa, Constância Ayres

## Abstract

Zika (ZIKV) and chikungunya (CHIKV) are arboviruses that cause infections in humans and can causeclinical complications, representing a worldwide public health problem. *Aedes aegypti* is the primary vector of these pathogens and *Culex quinquefasciatus*may be a potential ZIKV vector. This study aimed to evaluate fecundity, fertility, survival, longevity, and blood feeding activity in *Ae. aegypti* after exposure to ZIKV and CHIKV and, in *Cx. quinquefasciatus*exposed to ZIKV.Three colonies were evaluated: AeCamp(*Ae. aegypti -*field),RecL (*Ae. aegypti -* laboratory)and CqSLab (*Cx. quinquefasciatus -* laboratory). Seven to 10 days-old females from these colonies were exposed to artificial blood feeding with CHIKV or ZIKV. CHIKV caused reduction in fecundity and fertilityinthe natural population, AeCamp and reduction in survival and fertility in RecL.ZIKV impacted survival in RecL, fertility in AeCamp. and fecundity and fertility in CqSLab. Both viruses had no effect on blood feeding activity. These results show that CHIKV produces a higher biological cost in *Ae. aegypti*, compared to ZIKV, and ZIKV differently alters the biological performance in colonies of *Ae. aegypti* and *Cx. quinquefasciatus*. These results provide a better understanding over the processes of virus-vector interaction and can shed light on the complexity of arbovirus transmission.

## Introduction

Complex interactions between vectors and arboviruses determine the vector competence of a species. These interactions act in association with biotic and abiotic factors;therefore, these aspects of a species’ biological performance can determine its vectorial capacity [1]. Among these parameters, some are related to reproductive and survival capacity [2]. Studies have already demonstrated the influence of arbovirus infection on the reproductive capacity of females of different species [3–9].For example, the number of eggs produced in the first gonotrophic cycle indicates the total lay profile during the entire female life in mosquitoes[10]. Changes in longevity and survival and changes in blood meal activity have also been associated with arbovirus exposure or infection [4,7,8].

The Zika (ZIKV) and chikungunya (CHIKV) viruses are RNA arboviruses from the Flaviviridae and Togaviridae families, respectively, which have quickly spread in recent years to various parts of the world, including Brazil [11–13]. These pathogens are a major public health concern, as they cause infections in humans that can trigger neurological complications, such as the Guillain-Barré Syndrome [14]and the Congenital Zika Syndrome [15], as well as cardiac manifestations[16]and painful and disabling polyarthralgia [17].

Recife, a municipality located in the northeast region of Brazil,is known to have very favorable environmental conditions for mosquito reproduction and maintenance of high vector population densities. These conditions, when associated with vectorial capacity and presence of a susceptible human population, construct a scenario conducive to the rapid propagation of an arbovirus [18]. Additionally, the ability of the species *Ae. aegypti* and *Ae. albopictus*to transmit ZIKV [19–22] and CHIKV [23–25], is an important factor for the spread of these viruses in the Americas [12,26]. Also, it should be considered that other highly anthropophilic local species may be involved in arbovirus transmission as secondary vectors [27]. *Cx. quinquefasciatus*, for example, is an abundant species in many parts of the world, including Brazil. However, most studies on its role in ZIKV transmission were only published as of 2016 [28–34].

In general, the information available on different aspects of the vectorial capacity of *Aedes aegypti*is not sufficient to explain the dynamics of arbovirus transmission, especially after pathogenic infections, considering that viruses modulate several parameters of the biological performance of vectors, and the mechanisms responsible for this modulation are also not well understood [9]. Further studies addressing aspects of the vector capacity of species involved in arbovirus transmission can elucidate fundamental questions for the area, pointing out the real role of the vector in a given territory and in an epidemiological context. Thus, the present study aimed to evaluate the effect of exposure to ZIKV and CHIKV on fecundity; fertility; survival; longevity and blood meal activity in *Ae. aegypti* from the city of Recife (Pernambuco, Brazil) and in a laboratory colony, as well as the effect of ZIKV on *Cx. quinquefasciatus*, a laboratory colony, considering the same aspects, which are relevant for arbovirus transmission [4,7,35–37].

## Materials and Methods

### Study area

The study was carried out in Recife (8°03’S 34°52’W), capital of the state of Pernambuco, Brazil, located in the northeast region. In the dry season, rainfall is scarce in the region, but it can be intense in the rainy season (from April to July). The temperature and relative air humidity range from 22 to 32°C and from 70 to 90% at different times of the year [38]. Recife is an hyperendemic area, withmultiple arboviruses circulating simultaneously, and it has been considered as the epicenter of the firstZika outbreak in Brazil.

### Mosquito samples

Three different mosquito populations were used in the present study: two laboratory colonies (RecL and CqLab) and one natural population (AeCamp).The laboratory colony of *Ae. aegypti* (RecL) has beenmaintained without contact with any larvicide or adulticidesince 1996, when it was established from collections performedin the neighborhood ofGraças, in Recife[39]. The other *Ae.aegypti* population, called AeCamp, came from the field, and was established through egg collectionsperformed between 2017 and 2018, in 13 neighborhoods in Recife (Santo Amaro, Várzea, Afogados, DoisIrmãos, Apipucos, Monteiro, Ipsep, Boa Viagem, Nova Descoberta, Vasco da Gama, CidadeUniversitária and Mustardinha). Experiments with AeCamp were carried out with individuals of the F_2_ generation.As for the *Culex* species, we have used the*Cx. quinquefasciatus*laboratory colony, known as CqSLab, which was originated inthe municipalities of Ipojuca, Olinda and Jaboatão dos Guararapes (Metropolitan Region of Recife), and it has been maintained since 2011 [40].

Field collections were performed using BR-OVT traps [38], as well as larvae and pupae, through shelling made directly in the breeding sites. All colonies were kept in the insectary of the Entomology Department of the Instituto AggeuMagalhães (IAM-FIOCRUZ-PE), under controlled conditions of temperature (26 °C ± 1 °C), relative humidity (50 to 90%) and photoperiod (14:10 h - L/D). Larvae and pupaewere kept in breeding containers with potable water andwere fed cat food (friskies®). Adult mosquitoes were kept in aluminum mesh cages (50 × 40 cm) and were fed a10% sucrose solution *ad libitum* on a daily basis. For females, blood feeding was offered weekly, in an artificial feeder, using defibrinated rabbit blood (*Oryctolagus cuniculus)*.

### Virus strains

ZIKV (BRPE243/2015)(41) and CHIKV (PE480/2016) strains were obtained from patients residing in the State of Pernambuco, Brazil, during the 2015 and 2016 outbreaks, respectively. These viruses were kindly provided by Dr. MarliTenório (Virology Laboratory, FIOCRUZ – PE). Viruses were propagated in Vero cells and tittered as described in Guedes et al. [31].

### Artificial bloodmeal for virus infection

Two to three independent experiments were performed with ZIKV and CHIKV. For *Ae. aegypti*, each experiment was performed withtwo groups for each colony (RecL, AeCamp and CqSLab): exposed to the virus (E) and non-exposed (NE, the control group).Two hundred adult females were used for the control groups (NE) and 300, for the test groups; they were aged between 7 and 10 days of emergence. In all groups, females were starved for 24 hours prior to oral exposure to the virus.

Oral blood-feeding was provided with an artificial feeder comprised of a Petri dish and Parafilm®, with a mixture of cultures of Vero cells inoculated with virus and defibrinated rabbit blood, in a volume of 10 mL, in a 1:1 ratio, as described in Guedes et al. [31]. The stock virus dose used was 10^6^ and 10^9^ plaque forming units per ml (PFU/ml), for ZIKV and CHIKV, respectively. The negative controlwas a mixture of equal volume of virus-free cell culture and defibrinated rabbit blood. Approximately 0.5 ml of each mixture was aliquoted for further titration. Females were exposed to a blood meal for one hour.All females had three more blood meals devoid of virus, after exposure to the virus, with the objective of evaluating blood meal activity and keeping them in active gonadotropic cycles for a better evaluation of longevity. These blood meals were provided once a week for three consecutive weeks.

### Assessment of the biological parameters after exposure to viruses

For the analysis of the putative biological cost, groups were defined according to the results of exposure to the viruses. Thus, for ZIKV, groups were divided into three: not exposed (NE); exposed, but not infected (E) and exposed and infected (EI). For CHIKV, groups were divided into two: not exposed (NE) and exposed and infected (EI), considering that, for this virus, infection rates were higher than ZIKV (above 90%), which made the number of exposed individuals and not infected not enough for statistical analysis (representing 4 and 8% for RecL and AeCamp, respectively).

### Fecundity and fertility assessment

Approximately 24 hours after the blood meal, 50 engorged *Ae. aegypti* females from each experimental group were transferred to individual cages. On the 3^rd^day post-exposure (dpe), each cage receiveda container (measuring 3.14 cm^3^) to mimic an oviposition site that contained 30% grass infusion and, in the case of the experiments with *Ae. aegypti*,posture supports made of cardboard, measuring approximately 4 × 2 cm.

In the*Ae. aegypti*assays, cardboardswere changed twice a week, in three gonadotropic cycles. The collected eggs were counted through a stereomicroscope to determine the fecundity rate of each female. After 15 to 20 days of quiescence, the eggs were placed in recipients containing 2 mL of 30% grass infusionto stimulate the synchronous hatching of the larvae. These were counted to estimate the individual fertility rate of each female. Fecundity and fertility analyses were performed in the first gonadotropic cycle.

In the *Cx. quinquefasciatus*assays, the recipients were removed from the individual cages after each oviposition (once a week), in three gonotrophic cycles. The number of eggs present in each collected raft were determined with the aid of a magnifying glass, up to 24 hours after laying. Larvae hatching percentage was evaluated approximately 72 hours afteroviposition, because theeggs of this species do not go into quiescence.

### Survival and longevity assessment

Mortality notification was performed daily to assess survival and longevity. To detect viral infection, 10 females in each experimental group of *Ae. aegypti*, were collected at three time points: 7; 14 and 21 dpe(days post exposure). There was a total of 20 females from each group until the end of life. For these, FTA - *classic card Whatman*^®^ cards (cards), containing *Manuka honey blend* ® honey, were made available on the screens of the cages,from the 7th to the 14th dpe. All females contributed to survival, and longevity assessments up to the point at which they left the study. For the analysis of average lifetime (longevity), females collected on the 7th, 14th and 21st dpewere excluded.

*Cx. quinquefasciatus*females were only collected when death occurredthroughout the experiments. Additionally, FTA classic card(Whatman® card) containing Manuka honey blend® were made available on the screens of all cages, from the 7th to the 14th dpe to ensure viral detection, through saliva collection, since females had not been collected on 7th, 14th and 21st dpe, as performed for *Ae. aegypti*.

Survival and longevity analysis was also performed among *Ae. aegypti*females in relation to viral load, represented by the RNA copy number (CN), detected by RT-qPCR. A cut-off point was determined to define the groups (with the highest and lowest viral loads) based on the median CN values found for the infected females.

### Search for blood meal

To assess the influence of ZIKV exposureon the blood meal activity of *Ae. aegypti*, females were fed with blood devoid of virus at 7, 14 and 21 dpe. After each feeding event, the completely engorged females were selected and counted. For *Cx. quinquefasciatus*exposed to ZIKV and *Ae. aegypti* exposed to CHIKV, evaluations were carried out exclusively with the first post-exposure blood meal (7 dpe).

### RNA extraction and RT-qPCR

Females collected at 7, 14 and 21 dpe, as well as those that died during the study, were placed separately in 1.5 ml microtubes, containing 300 μl of a mosquito diluent and stored at −80 °C until RNA extraction and RT-qPCR, described in Guedes et al. [31], with some modifications. The *primers* used for detection of CHIKV and ZIKV viral particles are describedin Lanciotti et al. [42,43]. Virus detection was performed by quantitative RT-qPCR on a QuantStudio^®^ 5 Real-Time PCR system (Thermo Fisher Scientific, Waltham, MA, USA), according toconditions described in Guedes et al. [31]. Cycle threshold values (Ct) were used to estimate the amount of viral RNA, using the standard curve as a reference for each RT-qPCR assay, obtained through isolated transcripts from the ZIKV BRPE243/2015 and CHIKV PE480/2016 strains, as described in Kong et al. [44]. Negative controls for the feeding experiment and RT-qPCR consisted of mosquitoes fed with virus-free blood and water, respectively.

Whole mosquitoes were processed, except for *Ae. aegypti* collected at 7 dpe, whose abdomens and thorax were analyzed separately from the heads. To calculate the infection rate (IR), the number of positive females was divided by the total number of analyzed samples. To calculate the dissemination rates (DR), head samples or positive cards were used, divided by the total number of positive samples. Females and cards with Ct lower than 38 were considered as positive.

The FTA cards were placed in 1.5 mL tubes and stored at −80 °C until use. To prepare the inoculum, cards were cut using multipurpose scissors and placed in 1.5 mL tubes. Next, 400 μL of ultrapure water was added to each tube, following homogenization for 5 times for 10 seconds, with 5 minute-intervals. Finally, the cards were transferred to a 10 mL syringe and filtered to enable recovery of the eluate only. The prepared inoculums were stored at –80°C until RNA extraction and RT-qPCR were performed, following the same protocol used for detection of viral RNA in mosquitoes.

### Statistical analysis

A descriptive analysis was performed: the variables were presented through graphs, followed by the presentation of the confidence interval and the *p*-value. Normality assumptions were made by applying the Shapiro Wilk tests. To assess the differences in means for the independent variables, the T-Student test was used, when the assumptions of normality were met. Otherwise, the Mann-Whitney test was applied, and the medians were evaluated.Also, the Bartlett test was used to assess homogeneity. When the assumption of homogeneity was met, the ANOVA mean test was used with Tukey’spost hoc test;if not, the median was evaluated by applying the Kruskal-Wallis’ test, with the post hoc test for Fisher’s test. The survival curve was determined using the Kaplan-Meier plot. The Cox test was applied to assess survival, and the proportionalities were evaluated using the Schoenfeld test. Conclusions were taken at a significance level of 5%.Results of the analysis were obtained using the R Core Team (2022). R: A language and environment for statistical computing. R Foundation for Statistical Computing, Vienna, Austria. URL https://www.R-project.org/

## Results

### Infection and dissemination rates of Zika (ZIKV) and chikungunya (CHIKV) viruses

Infection and dissemination rates varied between the viruses and colonies evaluated, being higher in the experiments carried out with the chikungunya virus (IR of 96.00% for RecL and 93.00% for AeCamp). The dissemination rates of the same virus ranged between 100.00% and 94.00% for RecL and AeCamp, respectively (Table 1). With Zika virus, these values were lower in CqSLab (22.50% IR and 57.00% DR), in relation to the two colonies of *Ae. aegypti* (Table 1).

**Table 1.** Zika (ZIKV) and chikungunya (CHIKV) virus infection (IR) and dissemination (DR) rates detected in laboratory *Aedes aegypti* (RecL), field *Aedes aegypti* (AeCamp) and laboratory *Culex quinquefasciatus* (CqSLab) colonies

| Virus | Colonies | IR | DR |
| --- | --- | --- | --- |
| ZIKV | RecL | 75.00 | 90.00 |
|  | AeCamp | 68.00 | 70.00 |
|  | CqSLab | 22.50 | 57.00 |
| CHIKV | RecL | 96.00 | 100.00 |
|  | AeCamp | 93.00 | 94.00 |

The percentage of positive cards varied among the colonies and viruses analyzed. In mosquitoes infected with ZIKV,the percentage rate was 30% for RecL, and60% forAeCamp, while for CqSLabfrom the first experiment, the percentage was low: 14.28%. No evaluation was possible forthe cards from the second and third experiments carried out with this colonybecause contamination was detected in the samples. Surprisingly, no cards were found to be positive for CHIKV in the two colonies (RecL and AeCamp), and we have no explanation for this.

The ZIKV viral load (number of RNA copies per mL - CN) was significantly higher (*p* = 0.019) among females from the RecL colony who underwent a second blood meal in blood free of viral particles, at7 dpe). The mean number of RNA copies increased from 4.31E+11 among those who did not have a second meal, to 5.81E+11 among those who ingested blood at 7 dpe. As a function of time of life after infection, it was found that the CN was significantly higher among RecL females that had completed engorgement at 7 dpe(p = 0.008)and died between the 8th and 22nd dpe(Fig 1). Although it was not statistically significant, there was an increase in CNin the period between 8 and 22 days in AeCamp(Fig 1).

**Fig 1.**
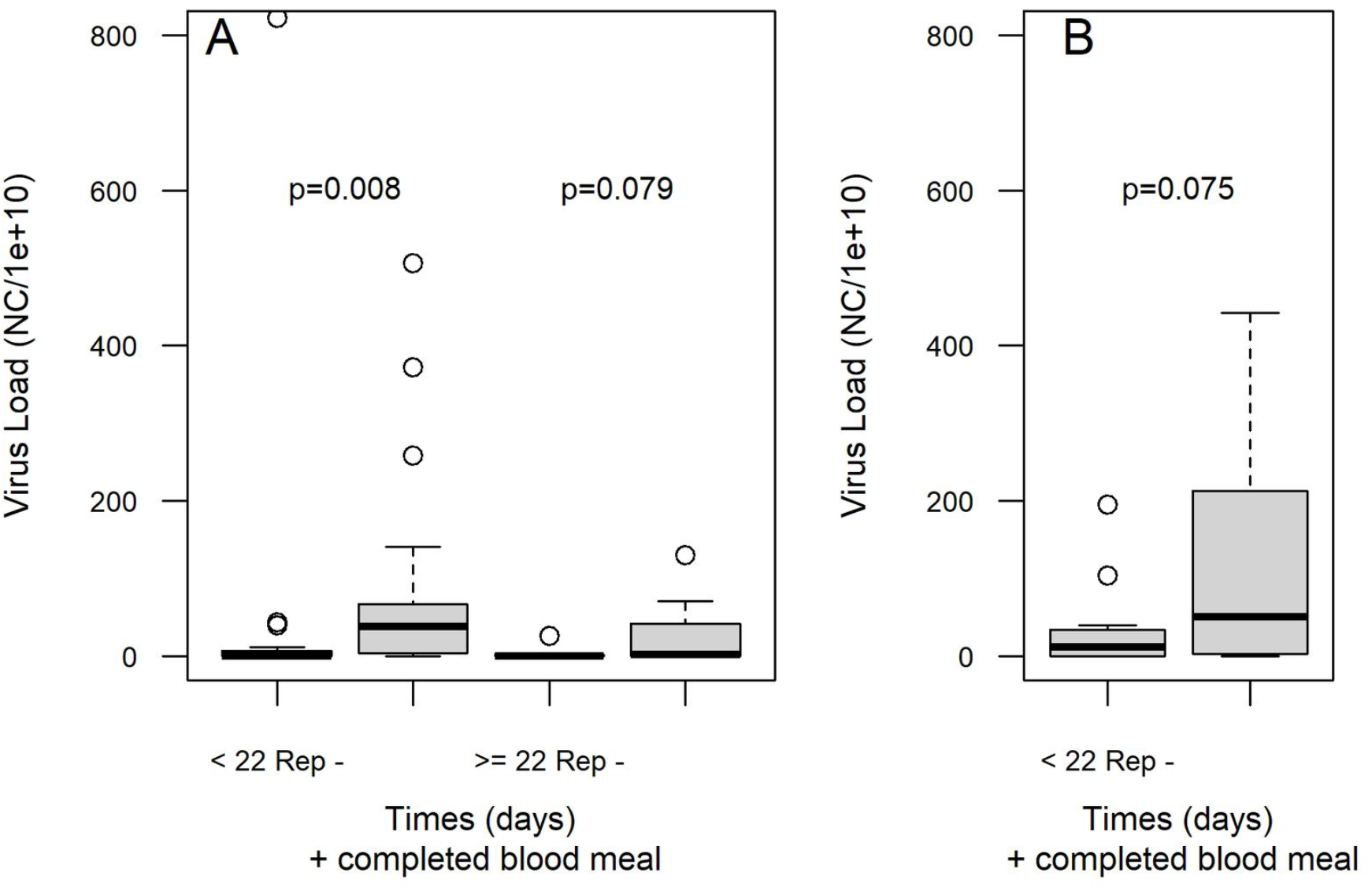
Viral load (number of RNA copies of ZIKV – NC, per mL, detected by RT-qPCR), in *Aedes aegypti* coloniesfrom the laboratory - RecL and field - AeCamp, infected with Zika virus (ZIKV), which completed blood meal on the 7th day post-exposure (dpe), in blood free of viral particles, as a function of post-exposure life span to the virus. A: RecL colony; B: AeCamp colony.Notes: < 22 Rep -: females that did not have a blood meal at 7 dpe post-exposure to ZIKV and died until the 21st dpe; >=22 Rep -: females that did not have a blood meal at 7 dpe post-exposure to ZIKV and died after the 21st dpe; < 22 Rep +: females who blood fed at 7 dpe to ZIKV and died until the 21 stdpe; >=22 Rep +: females that had a blood meal at 7 dpe post-exposure to ZIKV and died after the 21st dpe.

### Biological cost of infection with Zika (ZIKV) and chikungunya (CHIKV) viruses

In general, ZIKV and CHIKV had a significant impact on the parameters of vectorial capacity of the evaluated colonies. This impact negatively altered the reproductive capacity of females exposed to artificial oral infection by these arboviruses.

### Survival and Longevity

The analysis of the survival curve of the groups exposed to ZIKV, showed that the risk of death for females from RecL was about twice as high (E = 1.845: p = 0.014 and EI = 2.014: p = 0.003), compared to the control group (non-exposed - NE) (Fig 2A and S1-S3 Tables). Mean lifespans for the three groups did not differ significantly: 33.36; 31.04 and 26.23 days for NE, E and EI, respectively. However, for AeCamp, there was no significant difference in survival between the three groups analyzed, as the risk of death was 1,289 for group E and 1,212 for EI (Fig 2B and S1-S3 Tables). The average lifespan in AeCamp was also similar for the three groups: 34.19; 31.19 and 31.26 days, for NE, E and EI, respectively. Among *Cx. quinquefasciatus*females, survival (E = 1.300 and EI = 0.805) and longevity (34, 73; 31, 34 and 39, 50 days, for NE, E and EI, respectively) were not altered among females exposed to ZIKV (Fig 2C and S1-S3 Tables).

**Fig 2.**
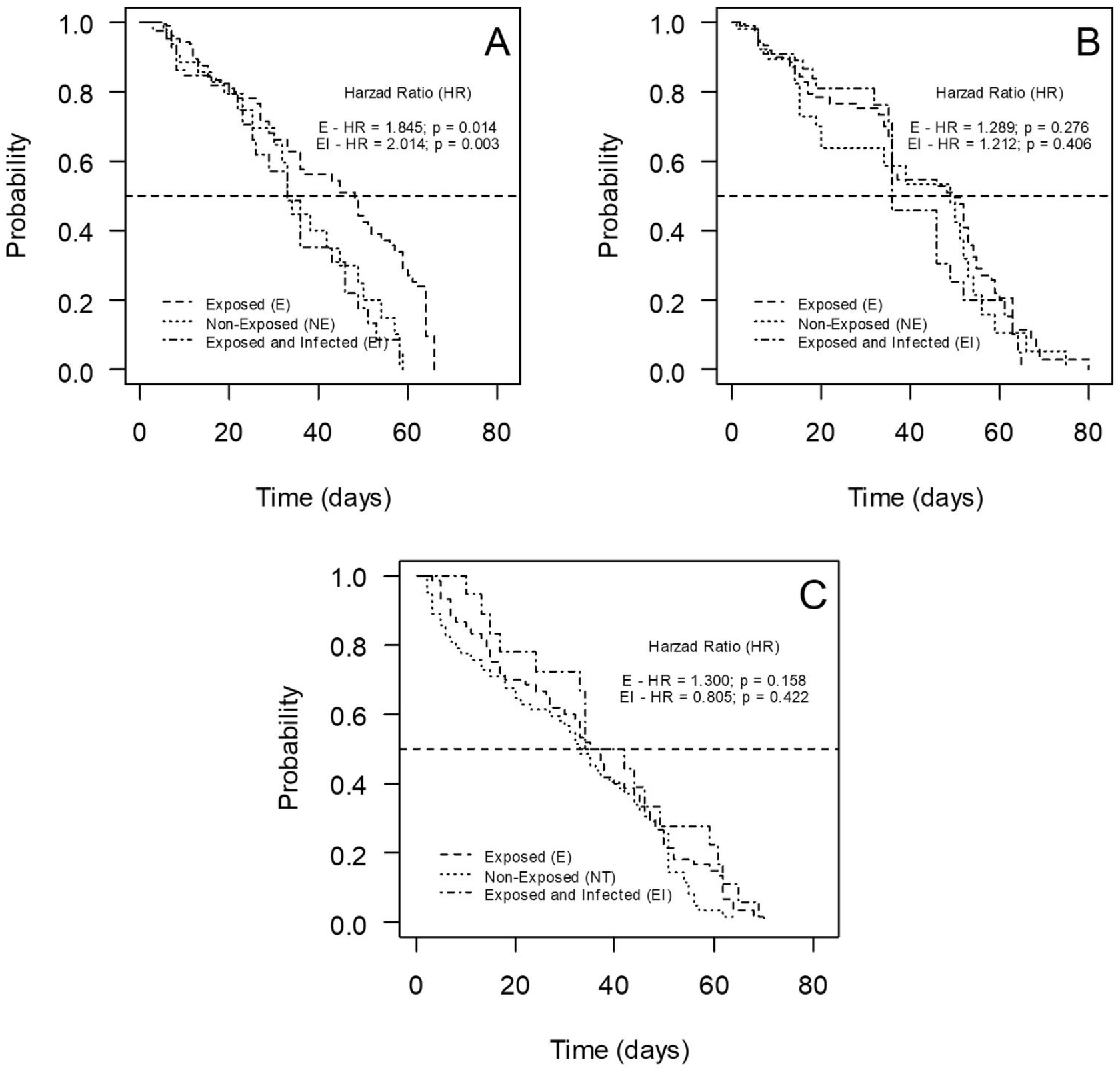
Survival curve of *Aedes aegypti* females– laboratory colony – RecL, over 66 days of observation; field colony - AeCamp, over 80 days of observation, and laboratory colony of *Culex quinquefasciatus*-CqSLab, over 70 days of observation, after exposure to Zika virus (ZIKV). A – RecL colony; B – AeCamp colony; C – CqSLab colony.

Likewise,the results found for the RecL colony, after exposure to CHIKV, showed an impact of infection on survival, with a higher risk of death (3.963) for the EI group (Fig3A and S1-S3 Tables); however, this difference appeared only between day zero and the 20th day of observation (p = 0.001), i.e., it was not found after the 21st day (Fig 3B). The longevity of RecL females was also reduced by CHIKV infection:38 and 17 days for NE and E, respectively (p = 0.002). On the other hand, in AeCamp, the survival curves showed no statistical difference between the two study groups (Fig 3C and S1-S3 Tables). Longevity was 37.5 days for the control group (non-exposed – NE) and 32 days for the EI group.

**Fig 3.**
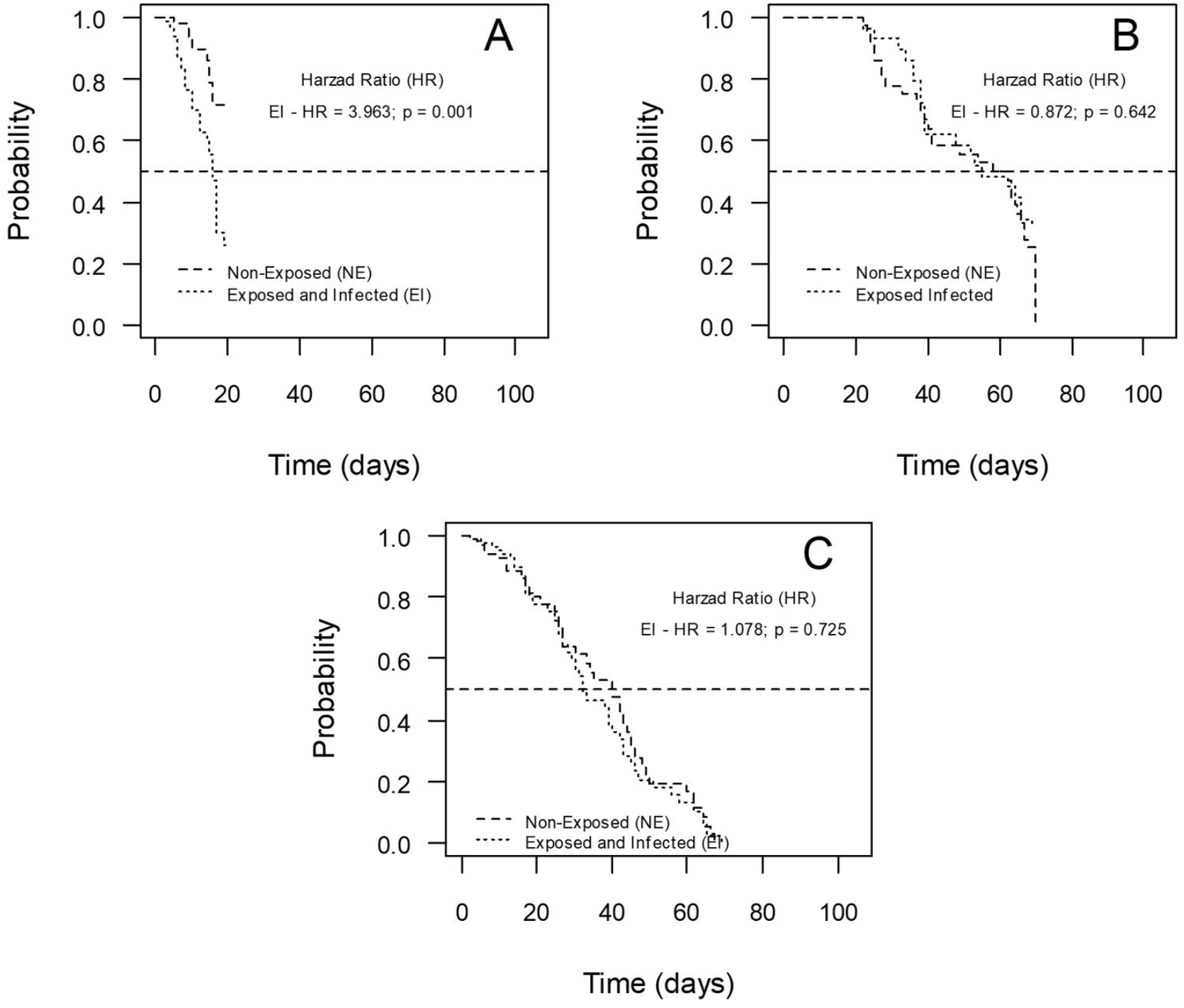
Survival curves: individuals from *Ae. aegypti* from laboratory - RecL, over 70 days and field - AeCamp, over 69 days of observation, after exposure to chikungunya virus (CHIKV). A - RecL colony - observation period less than or equal to 20 days; B - RecL colony - observation period longer than 20 days; C - AeCamp colony – observation period of 69 days.

### Fecundity and fertility in the first gonotrophic cycle

Fecundity (average number of eggs) and fertility (average percentage of larvae hatching) were not altered by exposure to ZIKV among *Ae. aegypti* from the RecL colony (Fig 4A), in the first gonotrophic cycle (number of eggs in the NE group = 87.83; E = 79.13; EI = 82.07 and average hatching percentage NE =67.36%;E= 66.31%; EI = 69.12%). In AeCamp, fecundity was also unaltered (number of eggs NE = 72.70; E: 70.40 and EI: 70.90) by exposure to ZIKV, although a significant reduction occurred in fertility (average percentage of hatching NE = 65.46%; E: 50.68% and EI: 49.38%) (Fig 4B). In *Cx. quinquefasciatus*, these parameters were not altered among infected females. However, fecundity and fertility were reduced in those exposed to the virus and whichdid not develop the infection (number of eggs = NE: 102.25; E: 76.90; EI: 92.67 and average hatching percentage NE = 74%;E = 58%; EI =69%) (Fig 4C).

**Fig 4.**
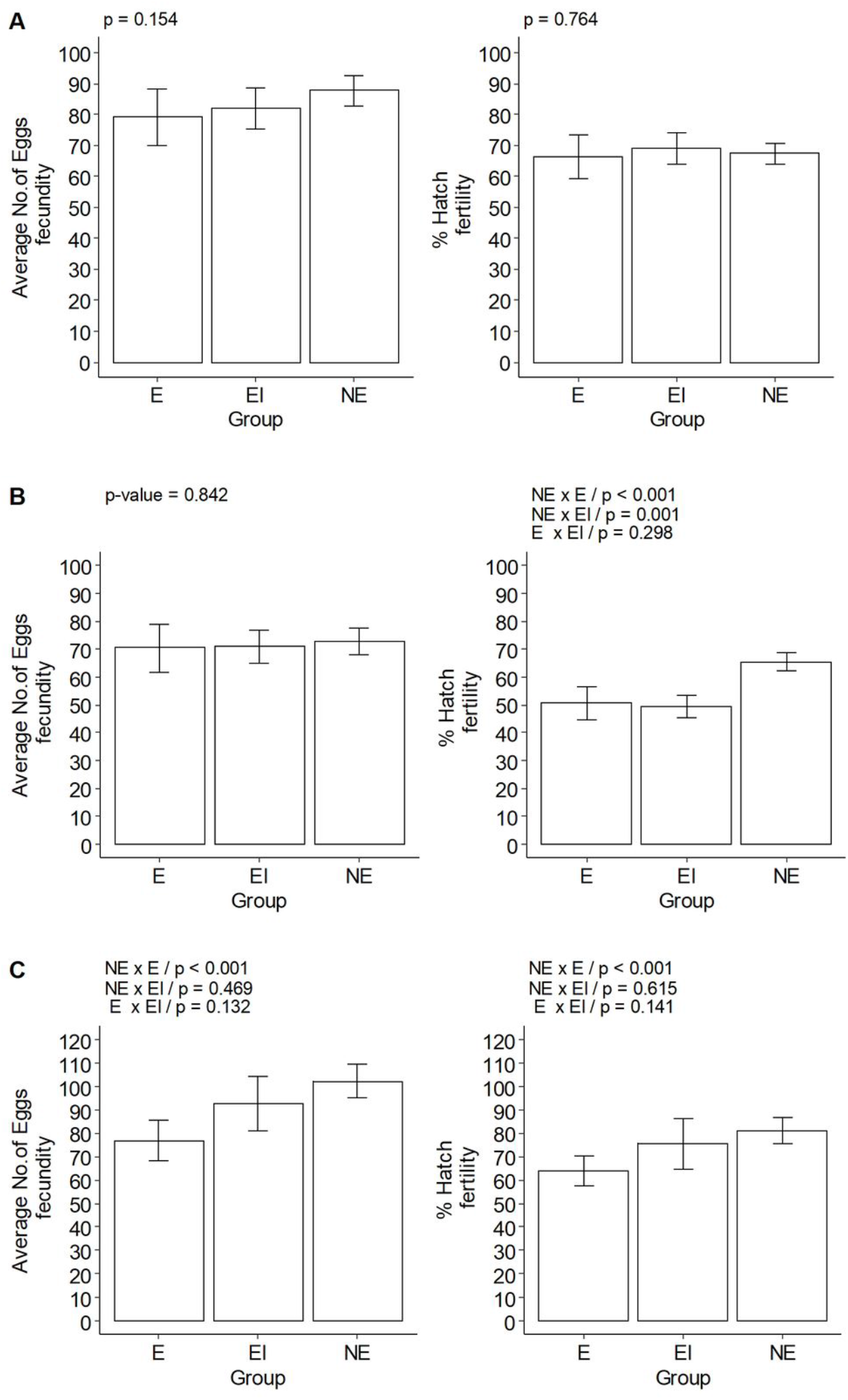
Average number of eggs laid and average percentage of larval hatching in the first gonadotropic cycle of *Aedes aegypti* females – laboratory colony – RecL and field colony – AeCamp, and of *Culex quinquefasciatus* females– laboratory colony – CqSLab, after exposure to Zika virus (ZIKV). Note: NE – Control (non-exposed); E - exposed; EI - exposed and infected. A: RecL colony; B: AeCamp colony; C:CqSLab colony.

The results show that the fecundity of females from the RecL colony was not significantly altered by the infection with CHIKV in the first gonadotropic cycle. However, the infection impacted fertility, with a reduction in the median percentage of hatching from 63.48 % in group NE to 40.67 % in group EI (Fig 5). Differently, in AeCamp, CHIKV had an impact on fecundity, reducing the median number of eggs from 48 in NE to 38 in group EI. The fertility of this colony was also altered by the infection, with a reduction in the median percentage from 57.50 to 37.50, between groups NE and EI, respectively (Fig 5).

**Fig 5.**
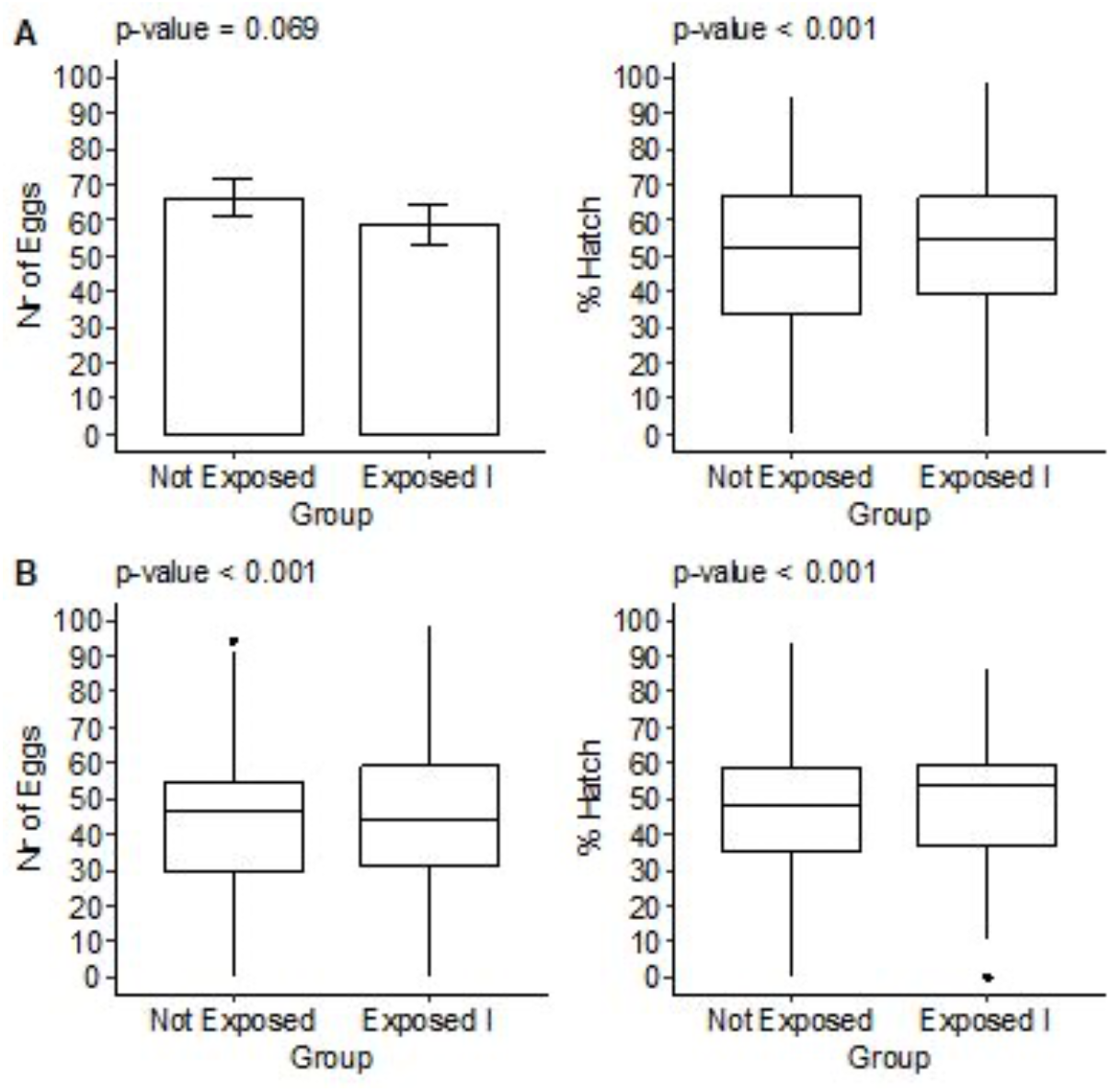
Mean or median number of eggs and median percentage of hatching of *Aedes aegypti* larvae– laboratory colony - RecL and field colony - AeCamp, in the first gonotrophic cycle, after exposure to chikungunya virus (CHIKV). Note: NE – Control (non-exposed); EI - exposed and infected. A: RecL colony; B: AeCamp colony.

## Blood Feeding Activity

The blood meal activity of RecL, AeCamp and CqSLab colonies was not altered by exposure to ZIKV and CHIKV, regarding the search for a second blood meal. Detailed numbers are shown inTable 2.

**Table 2.**
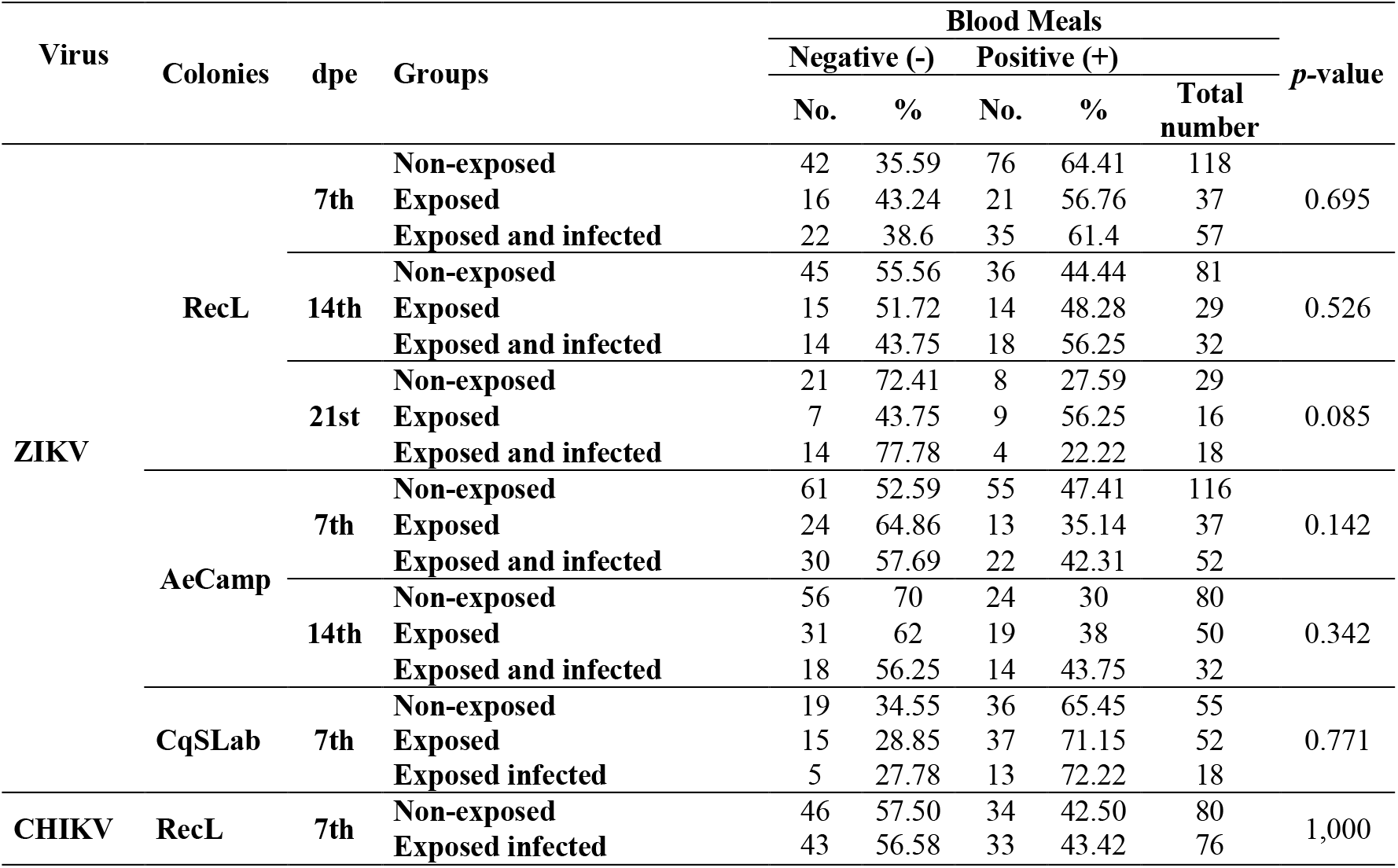
Number and percentage of females of *Aedes aegypti*, from colonies RecL and AeCamp, and of *Culex quinquefasciatus*,from colony CqSLab, which completed the blood meal in weeks following exposure to Zika virus (ZIKV) and from colonies RecL and AeCampexposed to chikungunya (CHIKV), in blood free of viral particles.

| Virus | Colonies | dpe | Groups | Blood Meals |  |  |  |  | *p*-value |
| --- | --- | --- | --- | --- | --- | --- | --- | --- | --- |
|  |  |  |  | Negative (-) |  | Positive (+) |  | Total number |  |
|  |  |  |  | No. | % | No. | % |  |  |
| ZIKV | RecL | 7th | Non-exposed | 42 | 35.59 | 76 | 64.41 | 118 | 0.695 |
|  |  |  | Exposed | 16 | 43.24 | 21 | 56.76 | 37 |  |
|  |  |  | Exposed and infected | 22 | 38.6 | 35 | 61.4 | 57 |  |
|  |  | 14th | Non-exposed | 45 | 55.56 | 36 | 44.44 | 81 | 0.526 |
|  |  |  | Exposed | 15 | 51.72 | 14 | 48.28 | 29 |  |
|  |  |  | Exposed and infected | 14 | 43.75 | 18 | 56.25 | 32 |  |
|  |  | 21st | Non-exposed | 21 | 72.41 | 8 | 27.59 | 29 | 0.085 |
|  |  |  | Exposed | 7 | 43.75 | 9 | 56.25 | 16 |  |
|  |  |  | Exposed and infected | 14 | 77.78 | 4 | 22.22 | 18 |  |
|  | AeCamp | 7th | Non-exposed | 61 | 52.59 | 55 | 47.41 | 116 | 0.142 |
|  |  |  | Exposed | 24 | 64.86 | 13 | 35.14 | 37 |  |
|  |  |  | Exposed and infected | 30 | 57.69 | 22 | 42.31 | 52 |  |
|  |  | 14th | Non-exposed | 56 | 70 | 24 | 30 | 80 | 0.342 |
|  |  |  | Exposed | 31 | 62 | 19 | 38 | 50 |  |
|  |  |  | Exposed and infected | 18 | 56.25 | 14 | 43.75 | 32 |  |
|  | CqSLab | 7th | Non-exposed | 19 | 34.55 | 36 | 65.45 | 55 | 0.771 |
|  |  |  | Exposed | 15 | 28.85 | 37 | 71.15 | 52 |  |
|  |  |  | Exposed infected | 5 | 27.78 | 13 | 72.22 | 18 |  |
| CHIKV | RecL | 7th | Non-exposed | 46 | 57.50 | 34 | 42.50 | 80 | 1,000 |
|  |  |  | Exposed infected | 43 | 56.58 | 33 | 43.42 | 76 |  |

## Discussion

Relevant aspects of the biological performance of vector mosquitoes can be altered by the process of pathogen infection, e.g., parameters involved in vectorial capacity. This event may lead to a consequent change in the pattern of occurrence of an epidemic in a given epidemiological context [45]. Studies have shown the biological cost of arbovirus infection, namely reduced survival, longevity, reproductive capacity, blood meal activity, among other impacts in mosquitoes. The different results found between them are explained by the direct relationship between the virus lineages and vectors involved [3,5–8,46]. Significant changes in reproductive capacity, for example, can limit the number of offspring of infected females and determine the transmission dynamics of an arbovirus. In general, this dynamic also occurs as a function of the age at which the vector acquires the infection, as well as survival and longevity [5,6,45].

In this sense, the two species investigated here, suffered a negative impact on biological performance, especially on reproductive capacity, after exposure to Zika (ZIKV) or chikungunya viruses (CHIKV), with a reduction in the number of individuals for the subsequent generation. This has evolutionary implications, once any trait that may offer an adaptive advantage for the mosquito’s defense against viral infection is unlikely to be selected for. However, the ability of a vector to transmit a pathogen is multifactorial and, therefore, isolated assessments in any of its parameters must be made with caution [45].

Our results showed that the natural population (AeCamp) did not suffer any impact on its survival or longevity caused by the infection of both viruses. This may suggest that infected mosquitoes can keep transmitting the virus for long periods in the environment. The interaction between vectors and viruses, genetically determined and triggered by the adaptive process in an environment, can result in a more efficient transmission dynamics, with less impact on the biological performance of *Ae. aegypti*[46].

On the other hand, for the *Ae. aegypti* colony, RecL, there was a significant reduction in survival when it was exposed and infected with both viruses. This result demonstrates a greater fragility of this colony, since the infection and dissemination rates for the two viruses were always higher when compared to the field population. It is known that the colonization process causes genetic diversity loss, which may impact on mosquito defense, metabolism, development, among other important traits for mosquito survival. Therefore, the results found in studies using laboratory colonies should always be interpreted with caution, before being extrapolated to what actually occurs in nature. The reduction in the survival rate and number of eggs laid by *Ae. aegypti* was associated with the process of adaptation to CHIKV infection, in a study developed by Sirisena, Kumar and Sunil [6]. The authors demonstrated that there is a negative regulation of genes involved in the egg laying pathway in infected females, through the analysis of transcript expression [6].

Previous studies have addressed cost on longevity [4] and survival [46]of *Ae. aegypti* populations infected by ZIKV. Petersen et al. [4], for example, used the ZIKV strain BRPE243/2015, the same strain used in the present study, to assess the reproductive capacity and longevity of *Ae. aegypti* collected in Rio de Janeiro, and they found a negative impact of the infection on the longevity of females. Thus, owing to genetic background and environmental factors, the viral strain used for infection in the laboratory should be considered, because especially in *Ae. aegypti*, the interaction between pathogen and vector can vary even between geographically close populations [46].

Similarly, the longevity of *Ae. aegypti* from Palm Beach County, Florida was also unaltered using the CHIK strain LR2006-OPY1. However, the same study suggests that physiological restrictions on the evolution of CHIKV infection in *Ae. albopictus* result in biological cost on this species, considering that they found a significant reduction in longevity. Additionally, the body titer of female *Ae. albopictus* infected with CHIKV and longevity were inversely correlated [47].

The number of eggs laid in the first gonotrophic cycle was not altered by exposure and infection with ZIKV in the two colonies of *Ae. aegypti*. On the other hand, the fertility of the field colony, AeCamp, was significantly reduced even among females exposed to ZIKV and that did not develop the infection. According to Li *et al*. [48], the ovaries of mosquitoes are affected by ZIKV on the second day after infection. Despite this fact, ZIKV did not reduce the fecundity of *Ae. aegypti*, as also found by Padilha et al. [3], Resck et al. [5] and Silveira et al. [46]. In contrast, a field population in Rio de Janeiro showed a reduction in fecundity. Interestingly, in the same study, one of the groups infected by ZIKV showed an increase in this parameter, between the first and third gonadotropic cycles [4].

The negative impact on AeCamp fertility suggested that the cost resulting from exposure to the virus was directed to the viability of the eggs, regardless of the establishment of the infection. Resck et al. [5] did not find an impact of ZIKV infection on the fertility of females from an *Ae. aegypti* from laboratory, corroborating the results reported in this paper forRecL. However, they differ from those found for the field colony. As demonstrated by Ciota et al. [49], the adaptive process is a determining factor for viral replication, in the case of successful infection, with subsequent increase in the immune response and consequent impact on the biological performance of the mosquito vector. This hypothesis explains the results regarding the fertility of AeCamp, which had contact with the strain used in these experiments during the Zika fever epidemic.

Parameters of reproductive capacity, especially fertility, showed a biological cost for infection with CHIKV, for both colonies, from the laboratory or field.Resck et al. [5] evaluated the reproductive capacity of *Ae. aegypti* after infection with CHIKV, and found a negative effect on fertility, but not on fecundity. A similar result was found with the CHIKV strain from Italy and *Ae. albopictus* from the same area. This alteration in fertility suggests that CHIKV can affect embryonic development and embryo survival [35]as described for other viruses [4,7]. In contrast, Carvalho-Leandro et al. [50]found no change in the reproductive capacity of *Ae. aegypti* infected with DENV, although the ovaries showed similar titers to other organs. The relevant effect of CHIKV infection on fertility, found in this study, suggests the need for further investigation, as this seems to be a critical point in the vector/parasite interaction mechanism that may guide the development of new vector control strategies.

For *Cx. quinquefasciatus*(CqSLab), survival and longevity were not affected by ZIKV exposure or infection. Styer, Meola and Kramer [7] found no difference in this aspect of vectorial capacity between *Cx. tarsalis* females, in a laboratory colony, after exposure to West Nile virus (WNV). The survival of *Cx. tarsalis*, however, was altered by Western equine encephalitis virus (WEEV) infection [51]. Exposure to ZIKV, but not infection, impacted the reproductive capacity of females, significantly reducing fecundity and fertility. For alaboratory colony of *Cx. pipiens*, pre-selected by continuous exposure to a WNV strain, fecundity was also altered; however, unlike what was found in the present study, infection, not just exposure, reduced this parameter [49]. The fecundity and fertility of *Cx. tarsalis* infected by WNV was also reduced in the infected groups [7]. Additionally, the same authors reported that the percentage of larvae hatching was higher in the exposed group (65.8%), than in the non-exposed (55.6%) and exposed and infected groups (42.5%).

The results analyzed for post-exposure blood feeding activity in an artificial feeder suggest that there is no effect of exposure or infection on the search for subsequent blood feeding in *Ae. aegypti* and *Cx. quinquefasciatus*, for both viruses and colonies studied here, considering the criterion evaluated (percentage of females that completed the blood meal within 30 minutes of exposure to the feeder). On the other hand, when evaluating the time spent by females of *Ae. aegypti* to complete engorgement, Sylvestre, Gandini and Maciel-de-Freitas [8]found that infected females spent more time compared to the non-exposed group. Additionally, a higher percentage of females of *Cx*. *tarsalis* infected with WNV had a blood meal on artificial feedings after infection, compared with unexposed females and exposed females that had not been infected. However, the same authors reported no significant difference in the amount of blood ingested between the three groups evaluated [7].

In general, the findings suggest that exposure to ZIKV and CHIKV significantly impact the reproductive capacity and longevity of the colonies evaluated. CHIKV had a greater impact on *Ae. aegypti*, in comparison to ZIKV, considering that the former reduced both parameters of the reproductive capacity of the field mosquitoes as well as the fecundity of RecL. The results also indicate that ZIKV differently impacted reproductive capacity when comparing *Ae. aegypti* and *Cx. quinquefasciatus* exposed and/or infected. Only exposure to ZIKV, but not infection, was enough to reduce the fecundity and fertility of *Cx. quinquefasciatus*females, suggesting that the triggering of defense mechanisms associated with the midgut barriers generates a biological cost for the species. Although *Cx. quinquefasciatus* showed much lower infection and dissemination rates than *Ae. aegypti*, this species is much more abundant in the environment in Recife; thus, its role in ZIKV transmission is not clear yet. In *Ae. aegypti*, ZIKV infection reduced the fertility of field females, but not their fecundity. However, this impact may have little relevance, considering that longevity, survival and search for blood source after exposure to the virus were not affected in the field population.

In the colony of *Ae. aegypti*, females infected with ZIKV, which had a second blood meal at 7 dpe, had a significantly higher number of RNA copies, compared to those that did not have a second meal. Similarly, Cui et al.[52] reported a three to four-fold increase in viral load among *Ae. aegypti* infected with DENV-4 after blood feeding at 5 dpe. However, owing to the limited number of females analyzed for this aspect, new experiments should be carried out, as they may allow a better statistical evaluation of this relationship, involving other variables to possibly confirm this result.

In summary, despite the significant reduction in some aspects of the biological performance of *Ae. aegypti*, for both study viruses, and for *Cx. quinquefasciatus*, with respect to ZIKV, it is suggested that the vectorial capacity of these species, supported by a successful global biological performance, close relationship with the host and availability of oviposition sites, poses a serious threat to public health, when associated with the susceptibility of the human population and the successful interaction with the pathogen. These results may, above all, increase the knowledge about the biological performance of mosquitoes in endemic and epidemic situations, in order to facilitate the definition and implementation of more effective methods of control of vector populations. Such knowledge can also trigger the development of future investigations, with a view to elucidating the several gaps that still exist on the various aspects of the vectorial capacity of mosquitoes.

## Supporting information

Support infomation (S1 table, S2 table and S3 table

## Acknowledgments

The authors are thankful to the insectary team for their support and for providing mosquitoes. To The Health Department of Recife. To Larissa Krokovsky for helpingwith RT-qPCR.

## Author Contributions

Conceptualization, M.C.,C.O., M.A.M.S., and C.A.; Methodology, M.C., E.H., M.S., R.A., R.B.; Formal Analysis, M.C., D.G., G.T., M.P.; Investigation, M.C.; Data Curation, M.C. and G.T.; Writing – Original Draft Preparation, M.C.; Writing – Review & Editing, all authors; Supervision, C.A.; Funding Acquisition, C.A. and M.P.

## Conflicts of Interest

The authors declare no conflict of interest.

### Ethical committee

This project was approved by the Research Ethics Committee of the InstitutoAggeuMagalhães-Fiocruz, Brazil (CAAE: 51012015.9.0000.5190).

## Supporting information

**S1 Table. Risk of death for females of***Ae. aegypti*, **RecL and AeCamp colonies, after exposure to Zika virus (ZIKV)**. Source: Fiocruz/PE. IAM’s Statistics and Geoprocessing Center.

**S2 Table. Risk of death for females from the laboratory***Aedes aegypti colony* **– RecL after exposure to chikungunya virus (CHIKV) during the first 20 days of observation**. Source: Fiocruz/PE. IAM’s Statistics and Geoprocessing Center.

**S3 Table. Risk of death for females from the field colony of***Aedes aegypti* **– AeCamp after exposure to chikungunya virus (CHIKV)**. Source: Fiocruz/PE. IAM’s Statistics and Geoprocessing Center.

## References

1. Kramer, L.D.; Ciota, A.T. Dissecting vectorial capacity for mosquito-borne viruses. CurrOpin Virol.2015, 15, 112–118. https://doi.org/10.1016/j.coviro.2015.10.003.

2. Barker, J.S.F. Defining fitness in natural and domesticated populations. In Adaptation and fitness in animal populations: evolutionary and breeding perspectives on genetic resource management. Van der Werf, J., Graser, H., Frankham, R., Gondro, C., Eds.; Springer: Heidelberg, Germany, 2009, pp. 3–14.

3. Padilha, K.P.; Resck, M.E.B.; Cunha, O.A.T.; Teles-de-Freitas, R.; Campos, S.S.; Sorgine, M.H.F.; Lourenço-de-Oliveira, R.; Farnesi, L.C.; Bruno, R.V. Zika infection decreases Aedes aegypti locomotor activity but does not influence egg production or viability. Mem Inst Oswaldo Cruz2018, 113, e180290. https://doi.org/10.1590/0074-02760180290.

4. Petersen, M.T.; Silveira, I.D.; Tátila-Ferreira, A.; David, M.R.; Chouin-Carneiro, T.; Van den Wouwer, L.; Maes, L.; Maciel-de-Freitas, R. The impact of the age of first blood meal and Zika virus infection on Aedes aegypti egg production and longevity. PLoS ONE, 2018, 13, e0200766. https://doi.org/10.1371/journal.pone.0200766.

5. Resck, M.E.B.; Padilha, K.D.; Cupolillo, A.P.; Talyuli, O.A.C.; Ferreira-de-Brito, A.; Lourenço-de-Oliveira, R.; Farnesi, L.C.; Bruno, R.V. Unlike Zika, Chikungunya virus interferes in the viability of Aedes aegypti eggs, regardless of females’ age. Sci Rep 2020, 10, 13642. https://doi.org/10.1038/s41598-020-70367-6.

6. Sirisena, P.D.N.N.; Kumar, A.; Sunil, S. Evaluation of Aedes aegypti (Diptera: Culicidae) life table attributes upon chikungunya virus replication reveals impact on egglaying pathways. J Med Entomol.2018, 55, 1580–1587. https://doi.org/10.1093/jme/tjy097.

7. Styer, L.M.; Meola, M.A; Kramer, L.D. West Nile virus infection decreases fecundity of Culex tarsalis females. J Med Entomol.2007, 44, 1074–1085. https://doi.org/10.1093/jmedent/44.6.1074.

8. Sylvestre, G.; Gandini, M.; Maciel-de-Freitas, R. Age-dependent effects of oral infection with dengue virus on Aedes aegypti (Diptera: Culicidae) feeding behavior, survival, oviposition success, and fecundity. PLoS ONE20138, e59933. https://doi.org/10.1371/journal.pone.0059933.

9. Javed, N.; Bhatti, A.; Paradkar, P.N. Advances in Understanding Vector Behavioural Traits after Infection. Pathogens2021, 10, 1376. https://doi.org/10.3390/pathogens10111376.

10. Suleman, M. Intraspecific variation in the reproductive capacity of Anopheles stephensi (Diptera: Culicidae). J Med Entomol.199027, 819–828. https://doi.org/10.1093/jmedent/27.5.819.

11. Faria, N.R.; Azevedo, R.S.S.; Kraemer, M.U.G.; Souza, R.; Cunha, M.S.; Hill, S.C.; Thézé, J.; Bonsall, M.B.; Bowden, T.A.; Rissanen, I. et al. Zika virus in the Americas: sarly epidemiological and genetic findings. Science2016, 352, 345–349. https://doi.org/10.1126/science.aaf5036.

12. Fernández-Salas, I.; Danis-Lozano, R.; Casas-Martínez, M.; Ulloa, A.; Bond, J.G.; Marina, C.F.; Lopez-Ordóñez, T.; Elizondo-Quiroga, A.; Torres-Monzón, J.A.; Díaz-González, E.E. et al. Historical inability to control Aedes aegypti as a main contributor of fast dispersal of chikungunya outbreaks in Latin America. Antiviral Res.2015, 124, 30–42. https://doi.org/10.1016/j.antiviral.2015.10.015.

13. Zanluca, C.; Melo, V.C.A.; Mosimann, A.L.P.; Santos, G.I.V.; Santos, C.N.D.; Luz, C. First report of autochthonous transmission of Zika virus in Brazil. Mem Inst Oswaldo Cruz2015, 110, 569–572. https://doi.org/10.1590/0074-02760150192.

14. Oehler, E.; Watrin, L.; Larre, P.; Leparc-Goffart, I.; Lastere, S.; Valour, F.; Baudouin, L.; Mallet, H.; Musso, D.; Ghawche, F. Zika virus infection complicated by Guillain-Barré syndrome: case report, French Polynesia. Euro Surveill.2014, 19, 20720. https://doi.org/10.2807/1560-7917.es2014.19.9.20720.

15. Albuquerque, M.F.P.M.; Souza, W.V.; Araújo, T.V.B.; Braga, M.C.; Miranda-Filho, D.B.; Ximenes, R.A.A.; Melo Filho, D.A.; Brito, C.A.A.; Valongueiro, S.; Melo, A.P.L. et al. The microcephaly epidemic and Zika virus: building knowledge in epidemiology. Cad Saúde Pública2018, 34, e00069018. https://doi.org/10.1590/0102-311X00069018.

16. Morrison, T.E. Reemergence of chikungunya virus. J Virol.2014, 88, 11644–11647. https://doi.org/10.1128/JVI.01432-14.

17. Madariaga, M.; Ticona, E.; Resurrecion, C. Chikungunya: bending over the Americas and the rest of the world. Braz J Infect Dis.2016, 20, 91–98. https://doi.org/10.1016/j.bjid.2015.10.004.

18. Boyer, S.; Calvez, E.; Chouin-Carneiro, T.; Diallo, D.; Failloux, A. An overview of mosquito vectors of Zika virus. Microbes Infec.2018, 20, 646–660. https://doi.org/10.1016/j.micinf.2018.01.006.

19. Dick, G.W.A.; Kitchen, S.F.; Haddow, A.J. Zika virus (I): isolations and serological specificity. Trans R Soc Trop Med Hyg.1952, 46, 509–520. https://doi.org/10.1016/0035-9203(52)90042-4.

20. Diallo, D.; Sall, A.A.; Diagne, C.T.; Faye, O.; Faye, O.; Ba, Y.; Hanley, K.A.; Buenemann, M.; Weaver, S.C; Diallo, M. Zika virus emergence in mosquitoes in Southeastern Senegal, 2011. PLoS One2014, 9, e109442. https://doi.org/10.1371/journal.pone.0109442.

21. Díaz-Quiñonez, J.A.; López-Martínez, I.; Torres-Longoria, B.; Vázquez-Pichardo, M.; Cruz-Ramírez, E.; Ramírez-González, J.E.; Ruiz-Matus, C.; Kuri-Morales, P. Evidence of the presence of the Zika virus in Mexico since early 2015. Virus Genes 2016, 52, 855–857.

22. Guerbois, M.; Fernandez-Salas, I.; Azar, S.R.; Danis-Lozano, R.; Alpuche-Aranda, C.M.; Leal, G.; Garcia-Malo, I.R.; Diaz-Gonzalez, E.E.; Casas-Martinez, M.; Rossi, S.L. et al. Outbreak of Zika virus infection, Chiapas State, Mexico, 2015, and first confirmed transmission by Aedes aegypti mosquitoes in the Americas. J Infect Dis.2016, 214, 1349–1356. https://doi.org/10.1093/infdis/jiw302.

23. Restrepo-Jaramillo, B.N. Infección por el virus del chikungunya. CES Medicina2014, 28, 313–323.

24. Enserink, M. Infectious diseases: massive outbreak draws fresh attention to little-known virus. Science2006, 311, 1085. https://doi.org/10.1126/science.311.5764.1085a.

25. Tsetsarkin, K.A.; Vanlandingham, D.L.; McGee, C.E.; Higgs, S. A single mutation in chikungunya virus affects vector specificity and epidemic potential. PLoS Pathog.2007, 3, 1895–1906. https://doi.org/10.1371/journal.ppat.0030201.

26. Chouin-Carneiro, T.; Vega-Rua, A.; Vazeille, M.; Yebakima, A.; Girod, R.; Goindin, D.; Dupont-Rouzeyrol, M.; Lourenço-de-Oliveira, R.; Failloux, A. Differential susceptibilities of Aedes aegypti and Aedes albopictus from the Americas to Zika virus. PLoSNeglTrop Dis.2016, 10, e0004543. https://doi.org/10.1371/journal.pntd.0004543.

27. Ayres, C.F.J. Identification of Zika virus vectors and implications for control. Lancet Infect Dis.2016, 16, 278–279. https://doi.org/10.1016/S1473-3099(16)00073-6.

28. Azar, S.R.; Diaz-Gonzalez, E.E.; Danis-Lonzano, R.; Fernandez-Salas, I.; Weaver, S.C. Naturally infected Aedes aegypti collected during a Zika virus outbreak have viral titres consistent with transmission. Emerg Microbes Infect.2019, 8, 242–244. https://doi.org/10.1080/22221751.2018.1561157.

29. Fernandes, R.S.; Campos, S.S.; Ferreira-de-Brito, A.; Miranda, R.M.; Silva, K.A.B.; Castro, M.G.; Raphael, L.M.S.; Brasil, P.; Failloux, A.; Bonaldo, M.C. et al. Culex quinquefasciatusfrom Rio de Janeiro isnotcompetenttotransmitthe local Zika virus. PLoSNeglTrop Dis.2016, 10, e0004993. https://doi.org/10.1371/journal.pntd.0004993.

30. Gomard, Y.; Lebon, C.; Mavingui, P.; Atyame, C.M. Contrasted transmission efficiency of Zika virus strains by mosquito species Aedes aegypti, Aedes albopictus and Culex quinquefasciatus from Reunion Island. Parasit Vectors2020, 13, 398. https://doi.org/10.1186/s13071-020-04267-z.

31. Guedes, D.R.; Paiva, M.H.; Donato, M.M.; Barbosa, P.P.; Krokovsky, L.; Rocha, S.W.S.; Saraiva, K.L.A.; Crespo, M.M.; Rezende, T.M.; Wallau, G.K. et al. Zika virus replication in the mosquito Culex quinquefasciatus in Brazil. Emerg Microbes Infect.2017, 6, e69. https://doi.org/10.1038/emi.2017.59.

32. Guo, X.; Li, C.; Deng, Y.; Xing, D.; Liu, Q.; Wu, Q.; Sun, A.; Dong, Y.; Cao, W.; Qin, C. et al. Culex pipiens quinquefasciatus: a potential vector totransmit Zika virus. EmergMicrobes Infect.2016, 5, e102. https://doi.org/10.1038/emi.2016.102.

33. Heitmann, A.; Jansen, S.; Lühken, R.; Leggewie, M.; Badusche, M.; Pluskota, B.; Becker, N.; Vapalahti, O.; Schmidt-Chanasit, J.; Tannich, E. Experimental transmission of Zika virus by mosquitoes from central Europe. Euro Surveill.2017, 22, 30437. https://doi.org/10.2807/1560-7917.ES.2017.22.2.30437.

34. Main, B.J.; Nicholson, J.; Winokur, O.C.; Steiner, C.; Riemersma, K.K.; Stuart, J.; Takeshita, R.; Krasnec, M.; Barker, C.M.; Coffey, L.L. Vector competence of Aedes aegypti, Culex tarsalis, and Culex quinquefasciatus from California for Zika virus. PLoSNeglTrop Dis.2018, 12, e0006524. https://doi.org/10.1371/journal.pntd.0006524.

35. Bellini, R.; Medici, A.; Calzolari, M.; Bonilauri, P.; Cavrini, F.; Sambri, V.; Angelini, P.; Dottori, M. Impact of chikungunya virus on Aedes albopictus females and possibility of vertical transmission using the actors of the 2007 outbreak in Italy. PLoS One2012, 7, e28360. https://doi.org/10.1371/journal.pone.0028360.

36. Zink, S.D.; Van Slyke, G.A.; Palumbo, M.J.; Kramer, L.D.; Ciota, A.T. Exposure to West Nile virus increases bacterial diversity and immune gene expression in Culex pipiens. Viruses 2015, 7, 5619–5631. https://doi.org/10.3390/v7102886.

37. Phumee, A.; Chompoosri, J.; Intayot, P.; Boonserm, R.; Boonyasuppayakorn, S.; Buathong, R.; Thavara, U,;Tawatsin, A.; Joyjinda, Y.; Wacharapluesadee, S. et al. Vertical transmission of Zika virus in Culex quinquefasciatus Say and Aedes aegypti (L.) mosquitoes. Sci Rep.2019, 9, 5257. https://doi.org/10.1038/s41598-019-41727-8.

38. Barbosa, R.M.R.; Regis, L.N. Monitoring temporal fluctuations of Culex quinquefasciatus using oviposition traps containing attractant and larvicide in an urban environment in Recife, Brazil. Mem Inst Oswaldo Cruz2011, 106, 451–455. https://doi.org/10.1590/S0074-02762011000400011.

39. Melo-Santos MAV de, Sanches EG, Jesus FJ de, Regis L. Evaluation of a new tablet formulation based on Bacillus thuringiensis sorovar. israelensis for larvicidal control of Aedes aegypti. Mem Inst Oswaldo Cruz. 2001 Aug;96:859–60.

40. Amorim LB, Helvecio E, de Oliveira CMF, Ayres CFJ. Susceptibility status of Culexquinquefasciatus (Diptera: Culicidae) populations to the chemical insecticide temephos in Pernambuco, Brazil. Pest Manag Sci. 2013 Dec;69(12):1307–14.

41. Donald et al., 2016 https://www.ncbi.nlm.nih.gov/pmc/articles/PMC5051680/

42. Lanciotti, R.S.; Kosoy, O.L.; Laven, J.J.; Panella, A.J.; Velez, J.O.; Lambert, A.J.; Campbell, G.L. Chikungunya virus in US travelers returning from India, 2006. Emerg Infect Dis.2007, 13, 764–767. https://doi.org/10.3201/eid1305.070015.

43. Lanciotti, R.S.; Kosoy, O.L.; Laven, J.J.; Velez, J.O.; Lambert, A.J.; Stanfield, S.M.; Duffy, M.R. Genetic and serologic properties of Zika virus associated with an epidemic, Yap State, Micronesia, 2007. Emerg Infect Dis.2008, 14, 1232–1239. https://doi.org/10.3201/eid1408.080287.

44. Kong, Y.Y.; Thay, C.H.; Tin, T.C.; Devi, S. Rapid detection, serotyping and quantification of dengue viruses by TaqMan real-time one-step RT-PCR. J Virol Methods 2006, 138, 123–130. https://doi.org/10.1016/j.jviromet.2006.08.003.

45. Mayton, E.H.; Tramonte, A.R.; Wearing, H.J.; Christofferson, R.C. Vector competence for Zika virus is not affected by mosquito age but vectorial capacity of Ae. aegypti is multifactorial and age-dependent. Parasit Vectors2020, 13, 310. https://doi.org/10.1186/s13071-020-04181-4.

46. Silveira, I.D.; Petersen, M.T.; Sylvestre, G.; Garcia, G.A.; David, M.R.; Pavan, M.G.; Maciel-de-Freitas, R. Reduction on Aedes aegypti lifespan but no effects on mosquito fecundity and oviposition success. Front Microbiol.2018, 9, 3011. https://doi.org/10.3389/fmicb.2018.03011.

47. Reiskind, M.H.; Westbrook, C.J.; Lounibos, L.P. Exposure to chikungunya virus and adult longevity in Aedes aegypti (L.) and Aedes albopictus (Skuse). J Vector Ecol.2010, 35, 61–68. https://doi.org/10.1111/j.1948-7134.2010.00029.x.

48. Li, C.; Guo, X.; Deng, Y.; Xing, D.; Sun, A; Liu, Q.; Wu, Q.; Dong, Y.; Zhang, Y.; Zhang, H. et al. Vector competence and transovarial transmission of two Aedes aegypti strains to Zika virus. Emerg Microbes Infect.2017, 6, e23. https://doi.org/10.1038/emi.2017.8.

49. Ciota, A.T.; Ehrbar, D.J.; Matacchiero, A.C.; Van Slyke, G.A.; Kramer, L.D. The evolution of virulence of West Nile virus in a mosquito vector: Implications for arbovirus adaptation and evolution. BMC Evol Biol.2013, 13, 71. https://doi.org/10.1186/1471-2148-13-71.

50. Carvalho-Leandro, D.; Ayres, C.F.J.; Guedes, D.R.D.; Suesdek, L.; Melo-Santos, M.A.V.; Oliveira, C.F.; Cordeiro, M.T.; Regis, L.N.; Marques, E.T.; Gil, L.H. et al. Immune transcript variations among Aedes aegypti populations with distinct susceptibility to dengue virus serotype 2. Acta Trop. 2012, 124, 113–119. https://doi.org/10.1016/j.actatropica.2012.07.006.

51. Mahmood, F. Reisen, W.K.; Chiles, R.E.; Fang, Y. Western equine encephalomyelitis virus infection affects the life table characteristics of Culex tarsalis (Diptera: Culicidae). J Med Entomol.2004, 41, 982–986. https://doi.org/10.1603/0022-2585-41.5.982.

52. Cui, Y.; Grant, D.G.; Lin, J.; Yu, X.; Franz, A.W.E. Zika virus dissemination from the midgut of Aedes aegypti is facilitated by bloodmeal-mediated structural modification of the midgut basal lamina. Viruses2019, 11, 1056. https://doi.org/10.3390/v11111056.

