## Supplementary material for "Exposure to Zika and chikungunya viruses impacts aspects of the vectorial capacity of *Aedes aegypti* and *Culex quinquefasciatus*": Support infomation (S1 table, S2 table and S3 table

### Supporting information

**S1 Table.** Risk of death for females of *Ae. aegypti*, RecL and AeCamp colonies, after exposure to Zika virus (ZIKV).

| Variables | RecL | | | | AeCamp | | | |
|---|---|---|---|---|---|---|---|---|
| | HR | 95% CI | | p value | HR | 95% CI | | p value |
| | | info | top | | | info | top | |
| Not exposed Group | 1,000 | | | | 1,000 | | | |
| Exposed Group | 1,845 | 1,130 | 3,013 | 0.014 | 1,289 | 0.8165 | 2,034 | 0.276 |
| Exposed infected Group | 2014 | 1,272 | 3,189 | 0.003 | 1,212 | 0.7696 | 1,910 | 0.406 |
| | Proportionality | | | | | | | |
| **Variables** | | Schoenfeld analysis | | **p value** | | Schoenfeld analysis | | **p value** |
| | | chic | | | | chic | | |
| Exposed | | 3.30 | | 0.070 | | 0.112 | | 0.738 |
| Exposed Infected | | 3.36 | | 0.067 | | 1,142 | | 0.285 |
| **GLOBAL** | | 4.87 | | 0.088 | | 1,582 | | 0.453 |

Source: Fiocruz/PE. IAM's Statistics and Geoprocessing Center.

**S2 Table.** Risk of death for females from the laboratory *Aedes aegypti colony* – RecL after exposure to chikungunya virus (CHIKV) during the first 20 days of observation.

| Variables | HR | survival | | p-value |
|---|---|---|---|---|
| | | 95% CI | | |
| | | info | top | |
| Not exposed Group | 1,000 | | | |
| Exposed Infected | 3,963 | 1.73 | 9.08 | 0.001 |
| **Variables** | | **Proportionality** | | **p-value** |
| | | **Schoenfeld analysis** | | |
| | | **chic** | | |
| Exposed Infected | | 0.0647 | | 0.799 |

Source: Fiocruz/PE. IAM's Statistics and Geoprocessing Center.

**S3 Table.** Risk of death for females from the field colony of *Aedes aegypti* – AeCamp after exposure to chikungunya virus (CHIKV).

| **Variables** | **HR** | **survival 95% CI info** | **survival 95% CI top** | **p-value** |
|---|---|---|---|---|
| Not exposed Group | 1,000 | | | |
| Exposed infected Group | 1.078 | 0.7092 | 1,638 | 0.725 |
| **Variables** | | **Proportionality Schoenfeld analysis chic** | | **p-value** |
| Exposed Infected | | 0.0048 | | 0.945 |

Source: Fiocruz/PE. IAM's Statistics and Geoprocessing Center.
